# Optimisation of cell fate determination for cultured muscle differentiation

**DOI:** 10.1101/2023.09.06.556523

**Authors:** Lea Melzener, Lieke Schaeken, Marion Fros, Tobias Messmer, Dhruv Raina, Annemarie Kiessling, Tessa van Haaften, Sergio Spaans, Arin Doǧan, Mark J. Post, Joshua E. Flack

## Abstract

Production of cultured meat requires defined medium formulations for the robust differentiation of myogenic cells into mature skeletal muscle fibers in vitro. Whilst such formulations can drive myogenic differentiation to an extent similar to serum-starvation based protocols, these cultures are invariably heterogeneous in nature, with a significant proportion of cells not participating in myofusion, limiting maturation of the muscle. Here, we use RNA sequencing to characterise this heterogeneity at single-nucleus resolution, identifying distinct cellular subpopulations, including proliferative cells that fail to exit the cell cycle, and ’reserve cells’ that do not commit to myogenic differentiation. We show that the ERK, NOTCH and RXR pathways act during the first stages of myogenic cell fate determination, and by targeting these pathways, cell cycle exit can be promoted whilst abrogating reserve cell formation. Under these improved culture conditions, fusion indices close to 100% can be robustly obtained in 2D culture. Finally, we demonstrate that this translates to higher levels of myotube formation and muscle protein accumulation in animal component-free bioartificial muscle constructs, providing proof of principle for the generation of highly differentiated cultured muscle with excellent mimicry to traditional muscle.

## Introduction

Cultured meat (also referred to as ’cultivated’ meat) is an innovative technology centred around the in vitro proliferation and differentiation of animal cells to yield edible food products for human consumption^1,2^. Interest in this field is driven primarily by the negative sustainability aspects concomitant with conventional meat production, ranging from greenhouse gas emissions to animal welfare and food security^3,4^. Despite this wealth of interest, the technology is at a nascent stage, with numerous challenges yet to be solved before mass commercialisation can be achieved^5–7^.

Most cultured meat bioprocesses aim to produce skeletal muscle tissue from agriculturally relevant species via the differentiation of stem cells, such as myogenic satellite cells (SCs), embryonic stem cells (ESCs) or induced pluripotent stem cells (iPSCs)^7–9^. A critical challenge for the design of such processes is the identification of culture conditions that drive robust, mature myogenic differentiation in vitro, affording accurate mimicry of the muscle component of conventional meat with respect to protein content, flavour, texture and mouthfeel^10^. Approaches involving genetic modification of cells, whilst powerful, are generally treated with caution due to regulatory and consumer acceptance concerns in many geographies^11^.

Myogenic differentiation requires the coordination of cell cycle exit with expression of a cascade of master transcriptional regulators^12,13^, including MyoD and myogenin. Although long studied, recreating this process in vitro is challenging, and skeletal muscle tissue engineering lags behind other tissue types (such as cardiac muscle). Recent years have seen important breakthroughs in this field, including the development of animal-free scaffolds and defined medium formulations that drive differentiation to levels comparable to traditional serum-starvation protocols for myogenesis^14,15^. Despite these advances, however, large populations of non-fusing cells are observed in the majority of myogenic culture conditions. Whilst this heterogeneity during differentiation is fairly well understood in vivo, where a small number of SCs adopt a non- differentiated, quiescent state to preserve the future ability of the muscle to regenerate^16,17^, the vast majority of cells do participate in myogenic fusion, in stark contrast to in vitro differentiation where fusion indices higher than 50% are unusual^15,18^. The non-differentiating cells are sometimes referred to as ’reserve cells’^19,20^, and though various pathways, including NOTCH signaling^21,22^, have been implicated in their specification, the mechanisms driving this cell fate decision in vitro remain unclear.

In this study, we used single nucleus mRNA sequencing (snRNA-seq^23,24^) to characterise heterogeneity in mono- and multinuclear cells during bovine myogenic differentiation in vitro, with the aim of understanding cell fate determination and using this knowledge to increase the proportion of cells that participate in differentiation and fusion, thus improving the quality of muscle tissue for cultured meat applications.

## Results

### Not all satellite cells fuse during myogenic differentiation (corresponding to Figure 1; Figure S1)

In order to reduce heterogeneity during muscle differentiation, we first sought to characterise this process in detail, by understanding the behaviour of cells that do not participate in myogenic fusion. We induced differentiation of primary bovine satellite cells (SCs), using the serum-free differentiation medium formulation we previously developed (hereafter referred to as ’SFDM v1’^15^; Fig. 1a). Expectedly, we found that a subset of SCs remain unfused even at the peak of differentiation (around 96 h), with a maximum desmin index (proportion of nuclei within desmin- stained areas) of around 60%, and significant numbers of nuclei visible in the areas between the myotubes (Figs. 1a, b). Beyond 96 h, a major phase of myotube detachment was observed, as evidenced by a major drop in desmin index, and the total and normalised desmin area (Figs. 1b - d). This phenomenon, in which myotubes detach from the substrate and eventually float in the culture medium, has been previously noted elsewhere^25,26^. Intriguingly, after near-complete myotube detachment, a ’second wave’ of differentiation was observed, with previously non-fusing cells forming small myotubes after 216 h (Figs. 1a - d), indicating that some SCs adopt a reversible non-myogenic cell fate decision during the early phase of differentiation.

**Figure 1:**
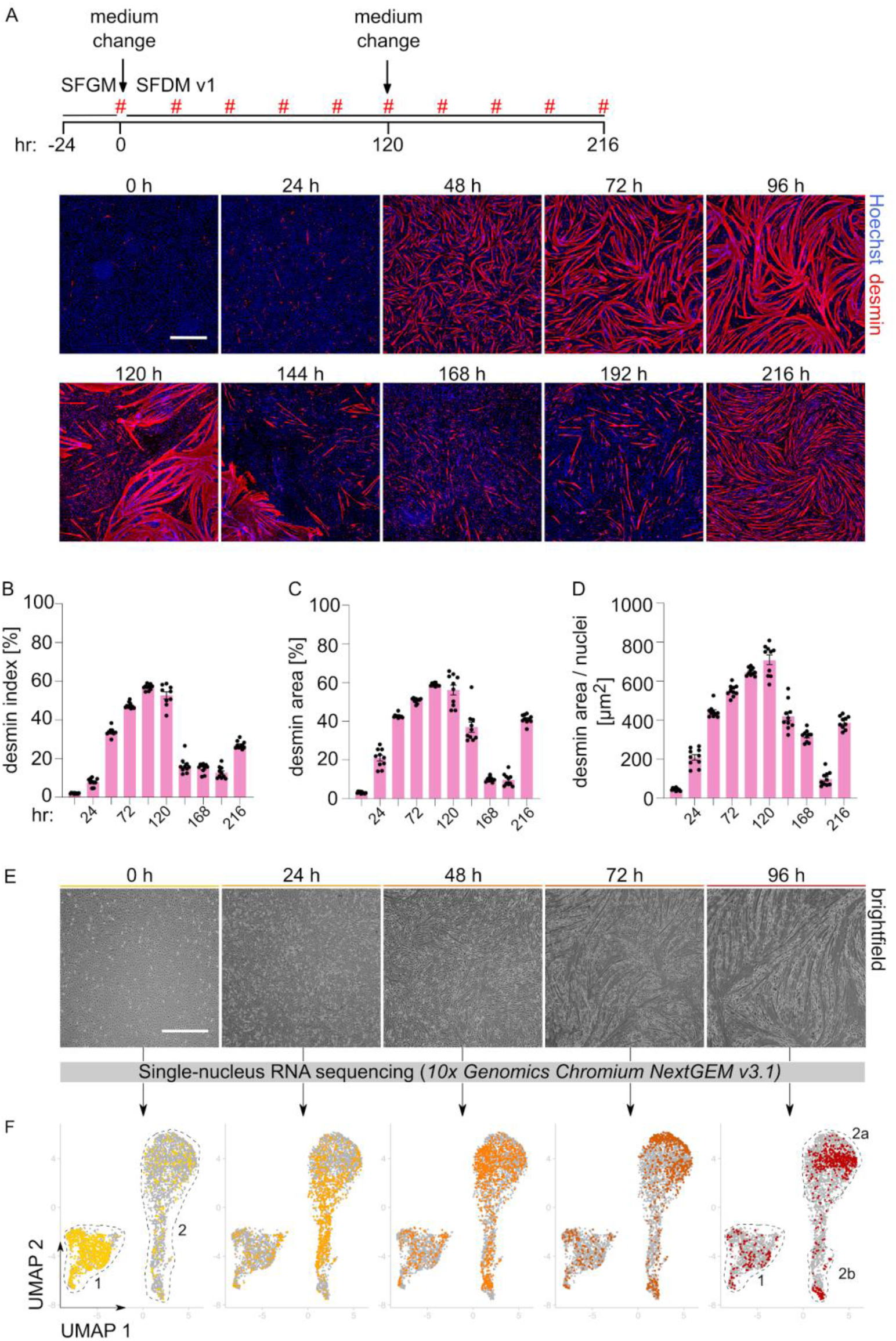
Only a subset of SCs fuse during myogenic differentiation. A: Fluorescence images of differentiating SCs to visualise myogenic fusion at timepoints indicated (#). Blue, Hoechst; red, desmin. Scale bar, 500 µm. B: Mean quantified desmin indices (proportion of nuclei within desmin stained areas) for images in A. Error bars indicate s.d., n = 10. C: Mean quantified desmin area (proportion of total image field that is desmin stained) for images in A. Error bars indicate s.d., n = 10. D: Mean quantified desmin area per nuclei for images in A. Error bars indicate s.d., n = 10. E: Representative brightfield images of differentiating SC cultures prior to nuclei harvest for snRNA-seq at 0, 24, 48, 72 and 96 h. Scale bar, 500 µm. F: Combined UMAP plots showing single nuclei from all five snRNA-seq timepoints (shown in E). Nuclei from each timepoint are coloured in the respective plot, all other nuclei are shown in grey. Nuclei clusters are annotated for clarity of explanation.

To further characterise the non-differentiating cells, we used a single-nucleus RNA sequencing (snRNA-seq^24,27^) approach to study transcriptional heterogeneity during the first 96 h of a standard differentiation culture in SFDM v1, containing a mixture of myotubes and mononuclear cells (Fig. 1e). A total of 3692 single-nuclei transcriptomes were analysed post-quality control (Supplementary Fig. 1), corresponding to 5 timepoints (0, 24, 48, 72 and 96 h). Clustering of these nuclei according to transcriptomic similarity revealed a number of distinct populations across the analysed timepoints (Fig. 1f), with marked transitions between timepoints also observable. At 0 h, two main populations were distinguishable (annotated ’1’ and ’2’), whilst by 24 h the relative abundance of the smaller of these populations (’2’) increased substantially. Intriguingly, this secondary population continued to evolve and subdivide over differentiation, with the nuclei thus appearing to fall into at least three distinct populations (’1’, ’2a’ and ’2b’) after 72 - 96 h (Fig. 1f).

### Single nuclei RNA sequencing identifies three distinct populations during differentiation (corresponding to Figure 2; Figures S2 & S3)

In order to identify the populations observed using snRNA-seq, we combined the nuclei transcriptomes from all timepoints into a single dataset (Fig. 2a), and analysed gene expression in and between the different nuclei clusters that were observed upon UMAP dimensionality reduction (based on the first 50 principal components). Based on shared nearest neighbour clustering, we were able to assign nuclei to three major populations, which we termed ’proliferating’, ’differentiating’, and ’reserve’ (Figs. 2a - c, Supplementary Figs. 2a - c). Only one of these populations (the ’differentiating’ nuclei) showed transcriptomic evidence of myogenic differentiation, as observed by expression of *MYOG* and enrichment of GO terms corresponding to genes involved in muscle contractility (Figs. 2b, c). Whilst the proliferating population was clearly marked by expression of known cell cycle genes, including *MKI67* and *CCND1*, the third population was slightly more enigmatic. Based on the elevated expression of *PAX7* (a known marker of quiescence) and absence of *MYOG,* we suspected that this likely represents a population of quiescent ’reserve’ satellite cells (Figs. 2b, d)^18^. This hypothesis was corroborated by calculating predicted cell cycle phases based on the snRNA-seq data (Supplementary Fig. 3a), as well as by analysis of highly expressed genes in a quiescent subpopulation we previously identified through scRNA-seq of SCs proliferating in vitro (Supplementary Figs. 3b, c)^28^. As well as the absence of an obvious myogenic signature, these reserve cells are easily distinguishable from the differentiating nuclei by their upregulation of a number of genes including *FTL*, *FTH1 and RPS2* (Fig. 2d). Interestingly, they also express *NOTCH2* and *HEYL* (a canonical NOTCH target gene), supporting a proposed negative feedback mechanism by which NOTCH signaling from differentiating cells (which themselves express *DLL1*) may help specify formation of reserve cells^22,29^.

**Figure 2:**
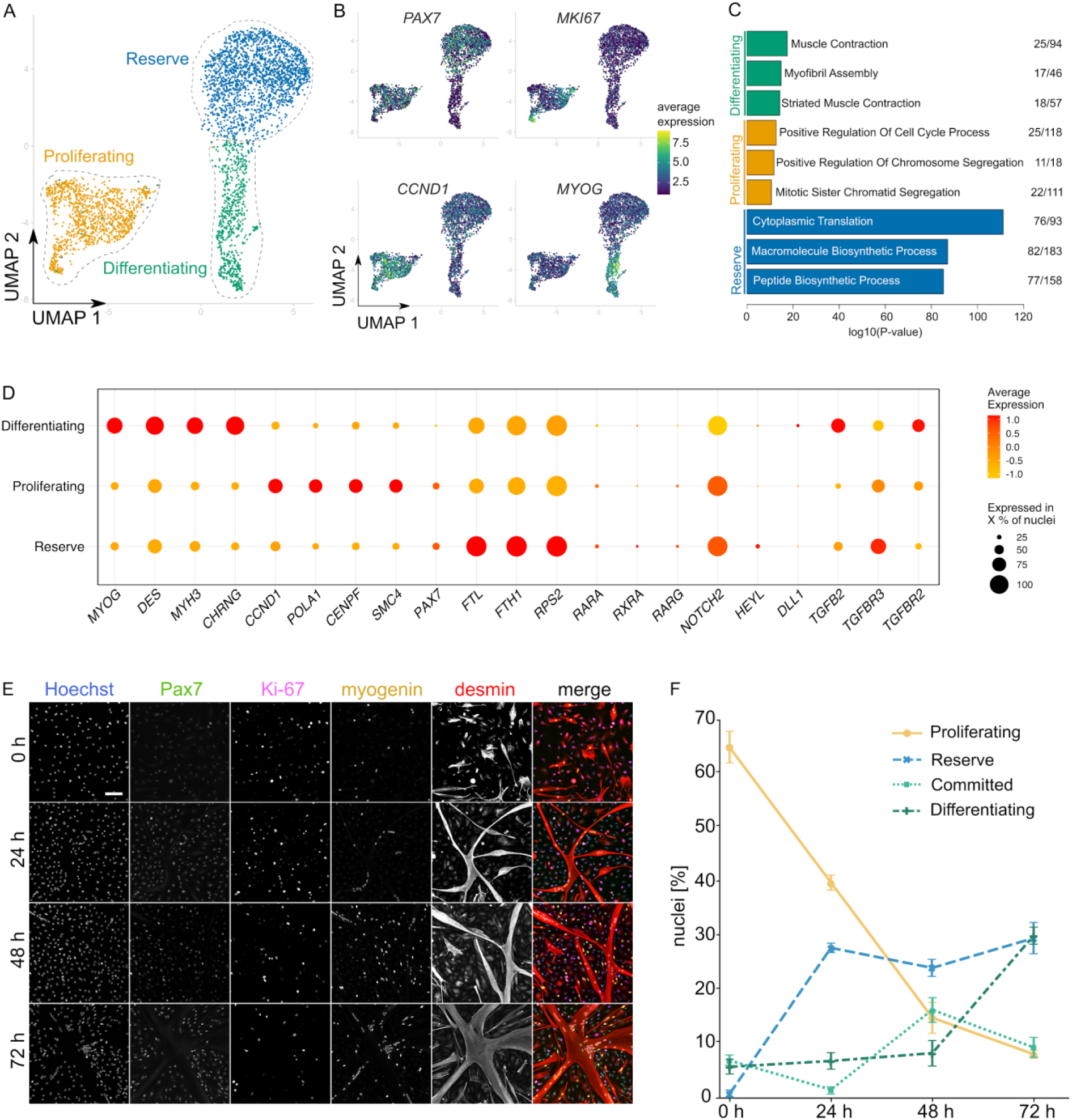
snRNAseq identifies committed, proliferating and reserve SC subpopulations. A: UMAP of single nuclei from all five timepoints, where nuclei are coloured and assigned according to identified clusters. B: UMAP as for A, coloured according to expression of indicated canonical marker gene(s) for each subpopulation. C: Significantly enriched GO terms corresponding to differentially expressed genes between the subpopulations identified in A. Numerals indicate proportion of genes for each GO term that are differentially expressed. D: Dotplot showing normalised expression of selected genes in each subpopulation (averaged across all nuclei); size of points indicates percentage of nuclei expressing the respective gene, colour indicates average expression level. E: Timecourse of fluorescence images for analysis of SC subpopulations (identified in A) during differentiation. Blue, Hoechst; green, Pax7; pink, Ki-67; yellow, myogenin; red, desmin. Scale bar, 100 µm. F: Quantification of SC subpopulations for images in E. Ki-67+, Proliferating; Pax7+/Ki-67-, Reserve; myogenin+/desmin-, Committed; myogenin+/desmin+, Myotube. Error bars indicate s.d., n = 3.

We also examined gene expression changes *within* these populations over the course of the experiment. Based on analysis of differentially expressed canonical marker genes for each population, we observed that whilst the proliferating nuclei generally show reduced expression of pro-proliferative genes (likely due to a reduction in pro-mitotic growth factor concentration in SFDM compared to SFGM), both the differentiating and reserve nuclei show increased expression of canonical markers over time (Supplementary Fig. 2b). This corroborates the idea that the differentiating and reserve populations diverge transcriptomically over time (as observed by UMAP; Fig. 1f), and suggests that cells exiting the cell cycle make a dynamic fate decision to adopt a differentiating or reserve cell fate.

In order to validate the nuclei clusters we observed in silico, we developed an immunofluorescent staining panel that allowed quantification of the snRNA-seq populations in vitro (Fig. 2e). SCs were differentiated in SFDM v1 for 72 h, and four nuclei populations quantified (Ki-67^+^, Proliferating; Pax7^+^/Ki-67^-^, Reserve; myogenin^+^/desmin^-^, Committed; and myogenin^+^/desmin^+^, Myotube), corroborating the heterogeneity observed by snRNA-seq (Fig. 2f). Expectedly, there was a significant drop in proliferating cells between 0, 24 and 48 h as cell cycle exit occurs, with a concomitant increase in differentiating (committed and myotube) nuclei. Notably, a substantial population of Pax7^+^ reserve cells formed after 24 h, and remained present at a relatively stable proportion throughout the timecourse. These heterogeneity dynamics were similar to those observed when quantifying the snRNA-seq data (Supplementary Fig. 2c), although some differences may arise from the efficiency of nuclei capture in our snRNA-seq protocol.

### Optimisation of serum-free differentiation medium (corresponding to Figure 3; Figures S4 & S5, Video S1)

We reasoned that manipulation of signaling pathways might alter the proportion of cells that adopt the proliferating, differentiating and reserve fates, and hence improve the overall extent of myogenic differentiation. Based on our transcriptomics data, combined with literature searches, we identified four major pathway manipulations (MAPK/ERK inhibition, RXR activation, NOTCH inhibition & TGF-β inhibition) that we hypothesised might increase the proportion of cells undergoing myogenic differentiation^22,30–32^. MAPK/ERK signaling is a well characterised pro-proliferative pathway in SCs, and along with RXR, has previously been exploited for improved myogenic differentiation^30,33^. Genes related to the RXR, NOTCH and TGF-β pathways, meanwhile, were upregulated in the ’reserve’ nuclei cluster of our snRNA-seq data (Fig. 2d).

For redundancy, we selected three small molecule compounds for each pathway manipulation and, after concentration optimisation, tested these as additives to SFDM v1 in a 2D myogenic differentiation assay (Fig. 3a, Supplementary Figs. 4a, b, Supplementary Table 3). For each pathway, at least one of the compounds showed an increase in myogenic differentiation as measured by desmin area (although for TGF-β inhibition, this was less significant). We therefore selected the most potent MEK/ERK inhibiting (MEKi; PD0325901, 1 µM), RXR activating (RXRa; ATRA, 500 nM) and NOTCH inhibiting (NOTCHi; DAPT, 5 µM) compounds for combinatorial testing. Whilst each of these compounds individually demonstrated a visible effect on myogenic differentiation after 48 and 72 h (Fig. 3b), desmin area measurements were not universally increased (Fig. 3c). However, the combination of all three molecules (hereafter referred to as ’SFDM v2’) showed significantly increased myotube formation at both timepoints, with almost all cells appearing to be involved in myofusion (Figs. 3b, c), as well as increased expression of muscle proteins, including myoglobin (Fig. 3d). Clear differences in differentiation kinetics were also observed using phase holographic live microscopy, including earlier myofusion and myotube detachment (Supplementary Video 1). We further demonstrated the relevance of the SFDM v2 formulation for cultured meat production, by comparing differentiation in SFDM v1 and v2 for SCs cultured in a 40 L stirred-tank bioreactor (Supplementary Figs. 5a, b), at different population doublings (PDs; Supplementary Fig. 5c), on a variety of different recombinant extracellular matrix (ECM) coatings (Supplementary Figs. 5d, e) and from three different donor animals (Supplementary Fig. 5f). In all cases, differentiation was significantly increased in SFDM v2 when compared to v1.

**Figure 3:**
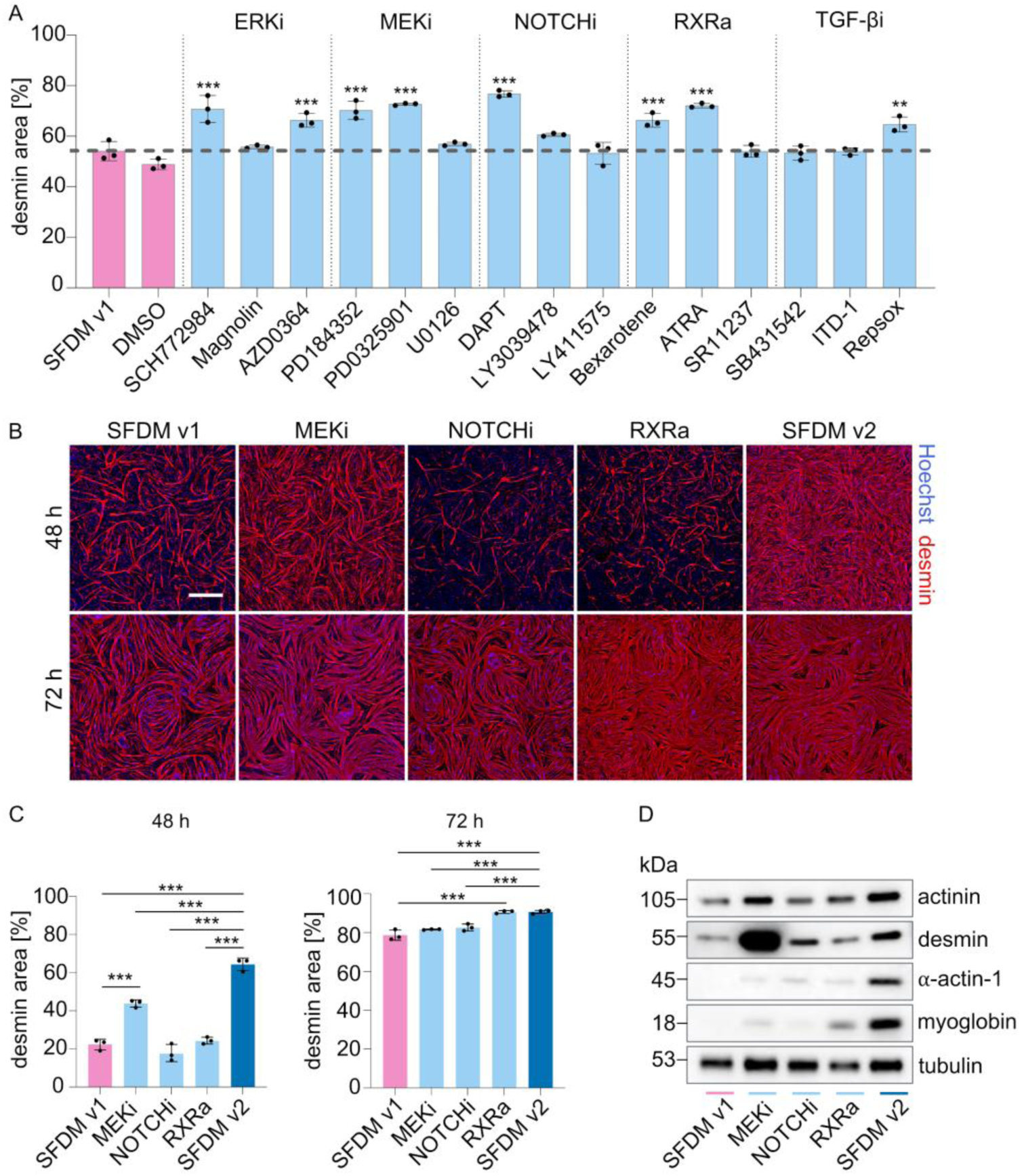
MEK/ERKi, NOTCHi and RXRa promote myogenic differentiation. A: Mean desmin areas after 72 h differentiation with indicated compounds (targeting MEK/ERK, RXR, NOTCH and TGF-β pathways). Error bars indicate s.d., n = 3. B: Representative fluorescence images of differentiation after indicated timeframe with three treatments showing significant effect in A (MEKi, PD0325901; NOTCHi, DAPT; RXRa, ATRA), and their combination (SFDM v2). Blue, Hoechst; red, desmin; Scale bar, 500 µm. C: Quantification of B, showing mean desmin areas after 48 (left) and 72 h (right) of myogenic differentiation with indicated compound treatment. Error bars show s.d., n = 3. D: Muscle-related protein expression, analysed by western blot after 72 h differentiation with the indicated treatment. *p<0.01, **p<0.001, ***p<0.0001.

### Differentiation-inducing molecules function by altering cell fate determination (corresponding to Figure 4; Figure S6)

We next sought to understand the mechanisms by which MEKi (PD0325901), NOTCHi (DAPT), RXRa (ATRA) and their combination (SFDM v2) lead to an increase in myogenic differentiation. Using the immunofluorescence staining panel previously designed (Fig. 2f), we examined the proportion of proliferating, differentiating and reserve nuclei in SFDM v1, each individual component, and SFDM v2 after 48 h of differentiation (Figs. 4a, b). For the individual components, the proportion of proliferating and (especially) reserve cells were significantly reduced when compared to SFDM v1, indicating treatment with these compounds can influence cell fate decisions during early differentiation. This effect was strongest for MEKi, although a synergistic effect was observed when all treatments were combined in SFDM v2, where no proliferating or reserve cells were observed whatsoever (Figs. 4a, b). Interestingly, whilst the SFDM v2 components, and MEKi in particular, do clearly promote cell cycle exit, treatment of SCs with other cell cycle-inhibiting compounds (abemaciclib, narazaciclib, palbociclib; targeting CDKs) had no significant effect on differentiation (Supplementary Figs. 6a, b), implying that MEKi acts to promote a specific cell exit towards a differentiating state.

**Figure 4:**
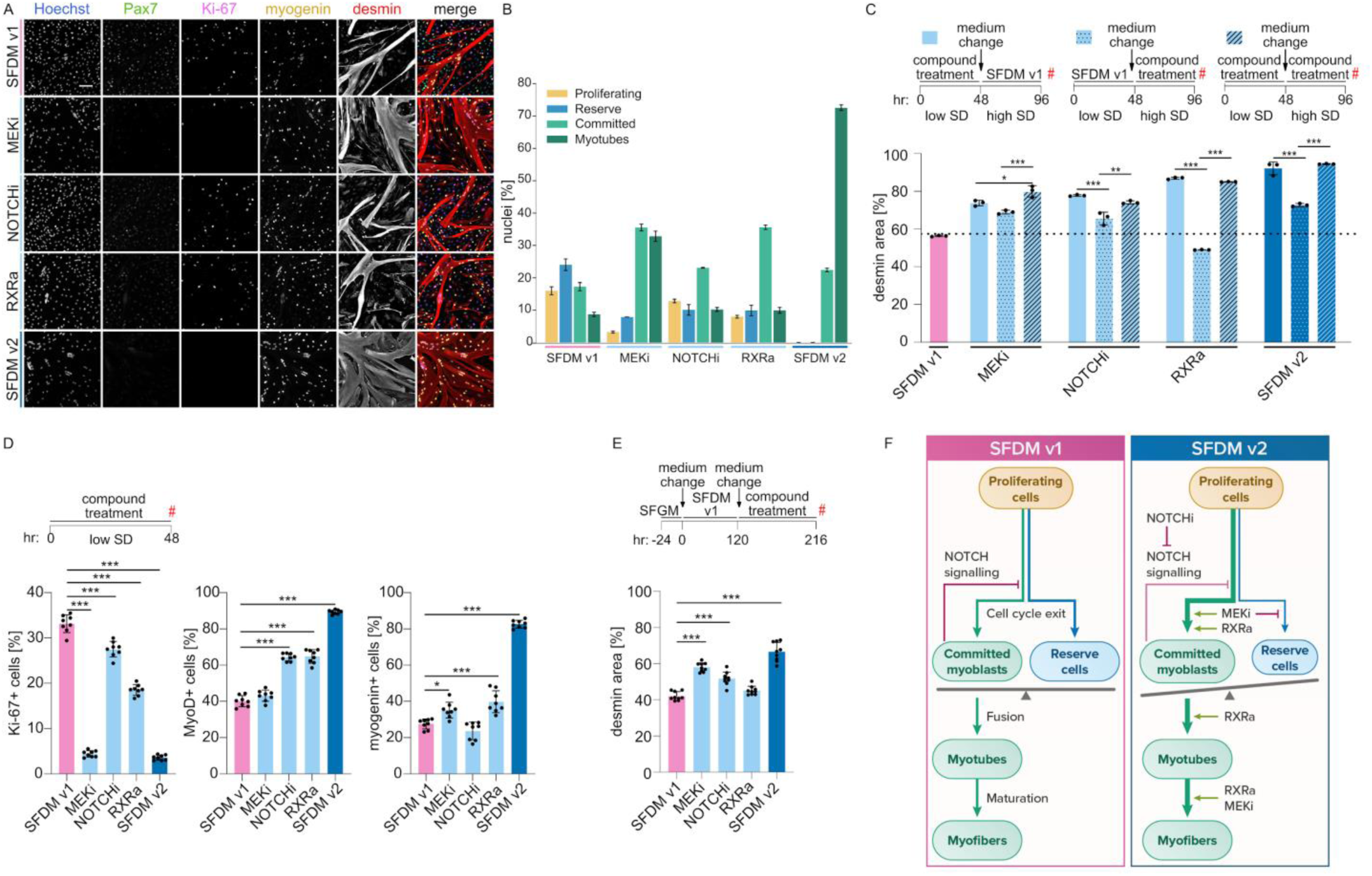
MEKi, NOTCHi and RXRa act during early commitment to myogenic differentiation. A: Representative fluorescence images after 48 h of differentiation for analysis of SC subpopulations during differentiation with indicated treatments. Blue, Hoechst; green, Pax7; pink, Ki-67; yellow, myogenin; red, desmin. Scale bar, 100 µm. B: Quantification of SC subpopulation images in A. Ki-67+, Proliferating; Pax7+/Ki-67-, Reserve; myogenin+/desmin-, Committed; myogenin+/desmin+, Myotube. Error bars indicate s.d., n = 3. C: Mean desmin areas at timepoint indicated (#) after compound treatment during only low seeding density (SD) phase, only high seeding density phase, or both. Error bars show s.d., n = 3. D: Quantification of Ki-67, MyoD and myogenin positivity after 48 h of indicated treatment at low seeding density. Error bars show s.d., n = 8. E: Mean desmin areas after compound treatment during the ’second wave’ of differentiation. Error bars show s.d., n = 9. F: Model proposing mechanistic basis for improved myogenic differentiation in SFDM v2. *p<0.01, **p<0.001, ***p<0.0001.

To investigate the hypothesis that SFDM v2 improves differentiation (at least in part) through alteration in the proportion of differentiating and reserve cells, we compared the effects of compound addition during a biphasic experiment. SCs were initially differentiated at a low seeding density (5×10^3^ cm^-2^; prohibitive to cell fusion), before reseeding at higher density (1×10^5^ cm^-2^) at which fusion is unimpeded by absence of cell-cell contacts (Fig. 4c). For all compounds, as well as for the combination, treatment during *only* the low seeding density phase lead to significantly greater differentiation than when cells were only treated at high density, suggestive of a role for these compounds in the early phase of cellular commitment to differentiation. We used immunofluorescence stainings for Ki-67, MyoD and myogenin to confirm that SFDM v2 treatment, even at low seeding density, leads to dramatically altered cell fate adoption. NOTCHi and RXRa alone significantly increased the proportion of MyoD+ cells, and demonstrated a highly synergistic effect with MEKi with respect to myogenin positivity (Fig. 4d, Supplementary Fig. 6c). Finally, we also investigated the potential for these compounds to drive reserve cell reprogramming, by studying their effect on reserve cells remaining in culture after myotube detachment. MEKi and NOTCHi treatment, as well as SFDM v2, significantly improved the second wave of differentiation, whilst RXRa had little effect (Fig. 4e, Supplementary Fig. 6d). Combined, these data are supportive of a model in which SFDM v2 promotes the myogenic differentiation of SCs by increasing the proportion of cells which initially adopt the differentiating cell fate, as opposed to the reserve cell state, as well as acting on downstream steps of muscle fusion and maturation (Fig. 4f)^30^.

### Differentiation inducers improve serum-free differentiation in bioartificial muscles (corresponding to Figure 5)

Accurate mimicry of traditional meat requires mature muscle differentiation in the context of a three dimensional tissue construct. We thus investigated these aspects of SFDM v2 induced myogenic differentiation. We first performed immunofluorescence staining for muscle proteins that are part of the contractile apparatus. SFDM v2 showed substantially increased expression of desmin, actinin and myosin, when compared to SFDM v1, indicative of more mature differentiation (Fig. 5a). Expectedly, this correlated with increased desmin, actinin and myosin indices (Figs. 5b - d). Close observation revealed increased levels of cross-striation in the SFDM v2 samples, particularly for actinin and myosin, further indication of mature differentiation (Fig. 5a).

**Figure 5:**
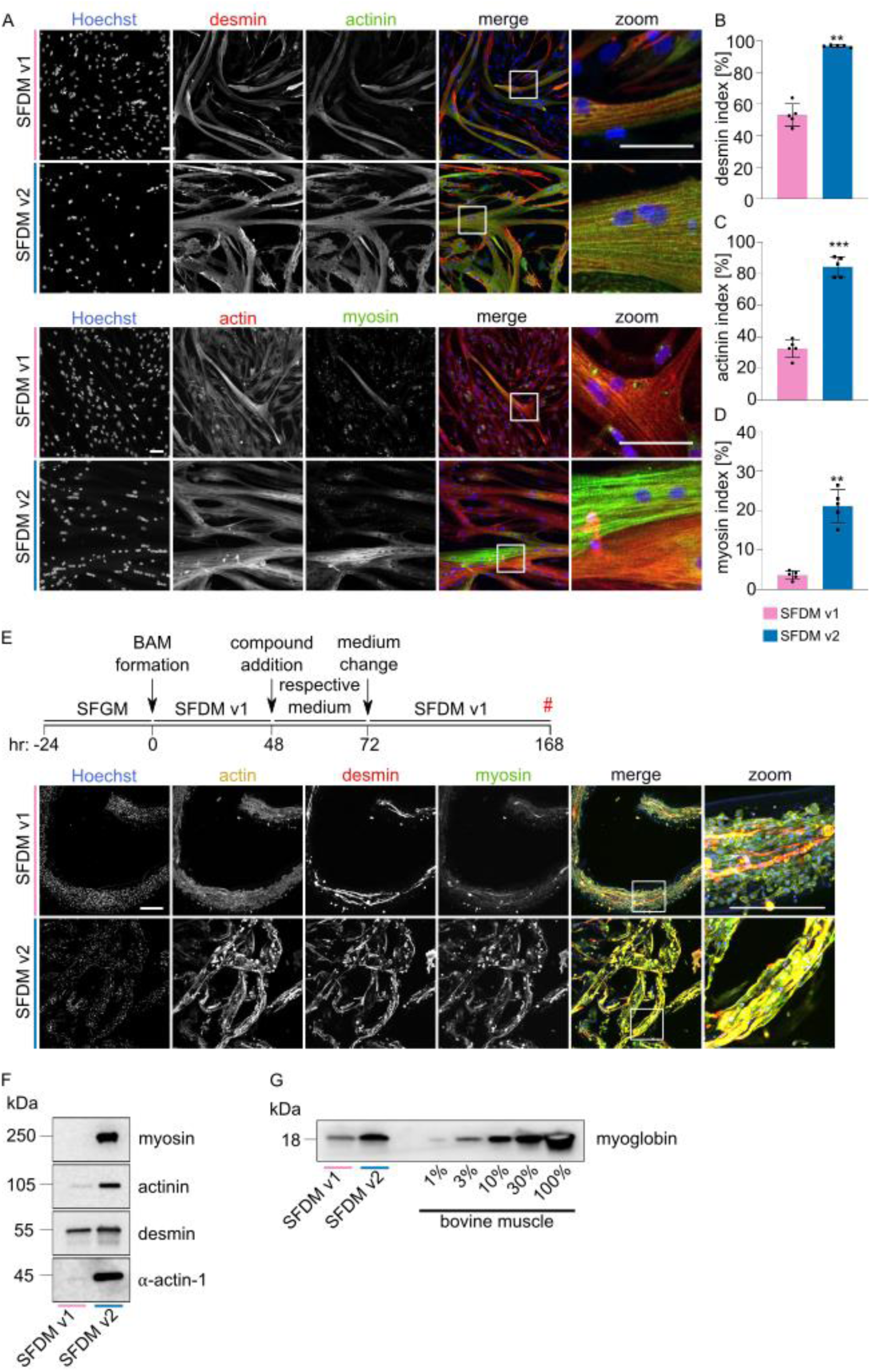
Inducers promote differentiation in animal-free bioartificial muscles. A: Representative fluorescence images after 72 h myogenic differentiation in SFDM v1 and SFDM v2 (indicated respectively) for two fluorescent staining panels. Blue, Hoechst; red, desmin/actin; green, actinin/myosin. Scale bar, 250 µm. B: Mean quantified desmin indices for images in A. Error bars indicate s.d., n = 5. C: As B, but actinin indices. Error bars indicate s.d., n = 5. D: As B, but myosin indices. Error bars indicate s.d., n = 5. E: Representative maximum-intensity projection fluorescence images after 7 days of 3D culture in an RGD-alginate-based animal-free bioartificial muscle system. Blue, Hoechst; yellow, actin; red, desmin; green, myosin. Lower panels show a zoom of the merged image. Scale bars, 200 µm. F: Muscle-related protein expression during 3D myogenic differentiation, analysed by western blot after 168 h differentiation with the indicated media. G: As F, for myoglobin expression and compared to a dilution series of bovine muscle tissue lysate. *p<0.01, **p<0.001, ***p<0.0001.

Finally, we tested the effect of SFDM v2 on myogenic differentiation in an animal-free, 3D bioartificial muscle (BAM) culture model, which would be suitable for upscaled and ethical cultured meat production. We used a self-assembling hydrogel system, in which RGD-functionalised alginate was extruded and gelated to create thin, SC-laden microfibers^34–36^. These structures were allowed to form in SFDM v1 for 48 h, during which time cell adhesion occurs, prior to treatment with SFDM v2 for a 24 h window (Fig. 5e). Longer treatment times with SFDM v2 lead to decreased viability of the constructs (data not shown). The extent of myotube formation was enhanced in constructs treated with SFDM v2 (Fig. 5e), whilst nuclei counts were clearly reduced, mirroring the results from 2D culture and suggesting that similar cell fate decisions operate during 3D differentiation. Expression of contractile muscle proteins, including myosin and actin, was also visibly enhanced, as corroborated by western blot, where levels of mature muscle proteins were clearly increased in SFDM v2 (Fig. 5f). We also compared myoglobin (a heme-containing protein critical for meat colour and quality^37^) expression against that in traditional meat, and found levels were substantially increased in SFDM v2 compared to v1, to around 30% of those found in bovine muscle tissue (Fig. 5g).

## Discussion

The production of cultured muscle tissue that accurately mimics conventional meat in appearance, texture, taste and nutritional value likely requires extensive and mature myogenic differentiation in vitro, a problem that is not yet solved. One complication is the failure of a proportion of myogenic cells to actually participate in differentiation and myofusion. We aimed to solve this bottleneck by first understanding the process of cell fate determination during in vitro myogenesis in depth, and then using these insights to improve upon our previous serum-free muscle differentiation medium formulation^15^.

Our initial observation of ’waves’ of differentiation during bovine SC myogenesis, whereby a second round of myotube formation occurs after the detachment of the ’primary’ myotubes, suggested a mechanism of reversible cell fate determination, in which not all cells differentiate despite having the capacity to do so (Fig. 1). Although such observations are not new^17,38^, the underlying pathways remain poorly understood. Our snRNA-seq approach, combined with immunofluorescence staining, revealed substantial heterogeneity during the early phase of differentiation (Fig. 2), pointing to two separate but interlinked issues; namely, that a subset of cells fail to efficiently exit the cell cycle in vivo, and of those that do, a large proportion enter a quiescent, non-differentiating state referred to as ’reserve cells’ (in line with previous literature^18^) that we were able to characterise transcriptomically for the first time. This phenomenon seems to occur at much higher rates in vitro than in the physiological setting^39^, but the precise extent to which these reserve cells are analogous to quiescent SCs in vivo remains difficult to understand. Indeed, the increased proportion of reserve cells observed in vitro likely reflects a lack of signaling from neighbouring cells or ECM that finely tune the number of SCs returning to quiescence in vivo^40^.

Based on analysis of our snRNA-seq data, together with literature research, we designed a screen for small molecule additives to improve our SFDM v1 formulation by increasing cell cycle exit and myogenic cell fate determination. Using myotube formation as a readout, we identified several different pathway manipulations that impacted differentiation, and which showed a strong additive effect (Fig. 3). MEK/ERKi and RXRa showed potent pro-myogenic effects, in corroboration with recent reports^30,41,42^. Interestingly, however, our data suggests that whilst there may be an effect on downstream steps of fusion, these pathways are also (or perhaps more) critical for cell fate specification during *early* differentiation (Fig. 4). Inhibition of NOTCH signaling likewise led to a reduction in reserve cell formation, synergistically with MEKi and RXRa. Differentiating cells express NOTCH ligands, including *DLL1*, whilst proliferating and reserve cells express *NOTCH2*, suggesting a negative feedback loop that creates a balance of differentiating and reserve cells^43^. In our system, the effect of TGF-βi was not as potent as other pathway manipulations, potentially because this is more critical for downstream fusion steps^31^, but further investigation of TGF-β (and related pathways, such as Wnt^44^) are nonetheless likely to yield additional improvements to SFDM v2. Furthermore, while it is clear that our pathway manipulations alter the balance of a critical cell fate decision, the precise timing of such a decision, relative to both initiation of differentiation, and the moment of cell cycle exit, remains unclear^45,46^. Whether reserve cells must reenter the cell cycle prior to differentiation also remains an open question, as does the requirement for a reduction in NOTCH signaling. Understanding these mechanisms will yield further opportunities to improve myogenic differentiation, and might be addressed with RNA staining or fluorescent reporter gene methodologies in future. A unified understanding of cell behaviour across the entire cultured meat bioprocess is important, and it is important to note that differentiation media must evolve in tandem with improved proliferation medium formulations.

Although for the purposes of cultured meat a single round of ’complete’ differentiation is likely optimal, a full understanding of the underlying biology is nonetheless invaluable, especially given the complexity of mimicking the entire muscle niche^47^. Whilst differentiation in our RGD-alginate hydrogel BAM system was strongly enhanced in SFDM v2 (Fig. 5), a small proportion of mononuclear cells were still present, indicating that achieving complete differentiation in 3D is particularly challenging. snRNA-seq of BAMs could be a useful method to understand how the additional stresses of these culture systems impact differentiation. Our protein analysis revealed that whilst myoglobin levels were substantially increased in SFDM v2, they are still below those of traditional beef, and use of hypoxic conditions or longer differentiation times might be helpful in this respect^48,49^. Interestingly, we found that reserve cells upregulate genes involved in iron storage and metabolism, including ferritin heavy (*FTH1*) and light (*FTL*) chains, but the relevance of this for myogenic differentiation is unclear^50^. Further improvements to 3D culture systems might also come in the form of additional functionalisation of alginate (to more accurately mimic the array of ECM proteins present in the muscle niche), addition of secondary components^51,52^, or the modulation of hydrogel stiffness^53^. We observed that treatment with SFDM v2 was optimal when initiated 48 h after BAM seeding, potentially because inhibition of MEK and/or NOTCH signaling can inhibit cell/substrate attachment during the initial BAM formation phase, but the timing of these pathway manipulations can likely be further tuned. Co-culture systems offer further intriguing possibilities for 3D differentiation^54^, as do the ’direct’ differentiation of cell aggregates^55^.

Another important question is whether our improved formulation would also work on a broader range of species, cell lines or cell types. Myogenesis is an ancient and well-conserved pathway^56^, and it is reasonable to suppose that the improved formulation could drive differentiation in other agriculturally relevant species, such as pigs or sheep, with minimal tweaking. Indeed, ERKi and RXRa have already been shown to be effective at improving differentiation in avian cells^30^, whilst NOTCHi has proven beneficial for fish myogenesis^57^. Whilst many proposed cultured meat bioprocesses utilise muscle-derived adult stem cells, either as unmodified primary cells, or as the basis for immortalised cell lines, a similar number propose the use of ESCs, iPSCs or non-muscle derived cell types, where the efficacy of this formulation would need to be tested^58^. If differentiation in these systems proceeds via a Pax7-expressing intermediate cell state, as has been observed in some systems^59,60^, many of the differentiation pathways implicated might again be involved, and our medium formulation might thus alleviate the need for inducible, transgenic approaches to muscle differentiation (which although biologically effective, leads to the undesirable categorisation of food products as genetically modified).

Cost reduction, safety and food regulation also represent critical considerations for myogenic differentiation medium formulations. The early-acting mechanism of SFDM v2 molecules might be advantageous in this respect, as treatment of cells during only the early days of a seven day (or longer) differentiation phase would allow for any residual levels to be ‘washed out’, although it remains to be seen how regulators will approach such issues. For some pathways, such as retinoic acid signaling (a derivative of Vitamin A), screening for food-safe inhibitors or agonists with similar myogenic effects is likely to be straightforward and potentially preferable^61^. Replacements for recombinantly-produced growth factor components of proliferation and differentiation media, with alternatives such as food-grade peptides or protein hydrolysates, might also help to reduce cost and regulatory scrutiny^62,63^.

Needless to say, numerous other scientific and non-scientific challenges remain to be solved before mass adoption of cultured meat can occur. However, the identification of conditions that can now drive highly efficient, complete myogenic differentiation in the absence of transgene expression represents an important step towards the efficient culture of meat.

## Methods

### Cell isolation and purification

Muscle-derived stem cells were isolated from bovine semimembranosus muscle immediately post-slaughter, using previously described protocols^28,64^. Briefly, muscle fibers were digested with collagenase (CLSAFA, Worthington; 1 h, 37 °C), followed by 100 μm and 40 μm cell filtration steps. Red blood cells were lysed with an Ammonium-Chloride-Potassium (ACK) lysis buffer (1 min, room temperature). Cells were plated in serum-free growth medium (SFGM, Supplementary Table 1) on fibronectin-coated (4 μg cm^-2^ bovine fibronectin, F1141, Sigma-Aldrich) tissue cultureware.

Satellite cells (SCs) were purified 72 h post-isolation by FACS as previously described^28^. Briefly, cells were stained with ITGA7-APC and ITGA5-PE (Supplementary Table 2) and sorted using a MACSQuant Tyto Cell Sorter (Miltenyi Biotec). Flow cytometry was performed regularly during SC proliferation using a MACSQuant10 Flow Analyzer (Miltenyi Biotec) to confirm cell purity.

### Cell culture

SCs were proliferated in defined serum-free growth medium (SFGM; Supplementary Table 1) on laminin 521 (0.5 μg cm^-2^, LN521-05, Biolamina) coated tissue culture vessels. After around 12 PDs (except where noted), SCs were differentiated on 0.5% Matrigel-coated cell culture vessels (except where noted) at a seeding density of 3.5×10^4^ cm^-2^. SCs were seeded in SFGM for 24 h, after which medium was exchanged to SFDM v1 (with respective compound addition where noted) to induce myogenic differentiation (Supplementary Tables 1, 3). Unless specifically noted, added compounds were included for the duration of the differentiation culture.

For bioreactor cell culture (Supplementary Fig. 4a), SCs were proliferated in SFGM on Cytodex- 1 microcarriers (Cytiva) in a 50c BioBLU single use vessel (Eppendorf; 37 °C, pH 7.2, 25 rpm) with a working volume of 40 L. Cells were grown to a final density of 2.9×10^6^ ml^-1^ prior to harvest for differentiation. Harvesting of a small sample was performed by incubation in 3X TrypLE (A1217702, Thermo Fisher Scientific; 30 mins, 37°C) to detach cells from microcarriers, and separated by 70 and 40 μm filtration steps prior to plating for differentiation as mentioned above.

### Immunofluorescent staining

Cells were fixed with 4% PFA, permeabilised with 0.5% Triton X-100 and blocked in 5% bovine serum albumin (BSA). Fixed cells were stained with rabbit α-desmin and respective secondary antibody (Supplementary Table 2) and Hoechst 33342 (Thermo Fisher Scientific), and imaged using an ImageXpress Pico Automated Cell Imaging System (Molecular Devices). Desmin index (proportion of nuclei within desmin stained areas), desmin area (proportion of total area that is desmin stained) and desmin area per nuclei were quantified using a customised protocol within the ImageXpress Pico analysis software. Except where otherwise noted in the figure legend, for each condition 3 separate wells were imaged, with each data point representing the analysis of a single well (itself the average of 9 separate images collected from within that well).

For immunofluorescence stainings for quantifying subpopulation heterogeneity (Figs. 2, 4) samples were stained with indicated antibodies, washed and incubated with respective secondary antibodies (Supplementary Table 2) and Hoechst 33342 (Thermo Fisher Scientific). Samples were imaged using a confocal microscope (TCS SP8, Leica Microsystems) with a 10x/1.00 objective. Confocal microscopy images were quantified in Fiji^65^ using homemade scripts. Differentiating nuclei were subdivided into ‘Committed’ and ‘Myotube’ populations in the immunofluorescence data where spatial information was retained; nuclei were assigned to the ’Myotube’ population (Figs. 2f, 4b) when they were located within a contiguous desmin stained area containing at least 2 nuclei. Images from three separate wells were analysed for each condition.

### Single nuclei RNA-sequencing

#### Cell lysis and nuclei harvesting

For each timepoint, SCs from two different animals were seeded at 5×10^4^ cm^-2^ on subsequent days and differentiated in SFDM for 96 h, 72 h, 48 h, 24 h or 0 h, resulting in a total of 10 samples with unique genotypes. On the day of harvest, nuclei were harvested by adapting a previously published protocol^24^. Cells were kept on an ice-cooled platform, washed with 1% BSA + 0.25 M Sucrose and lysed with 2.5% Triton-X100 for 10 min. The lysate was collected using a cell scraper, diluted with 1 % BSA + 0.25 M Sucrose, filtered with a 100 µm strainer and centrifuged to a crude pellet (3000 × g, 10 min). Nuclei pellets for each genotype were resuspended in Injection-buffer (1% BSA/RNase-free PBS + RNAse inhibitor), filtered with a 40 µm filter and quantified using a Countess 3 cell counter (Thermo Fisher) prior to injection.

#### Library preparation and sequencing

Quantified nuclei from all timepoints were mixed in an equal ratio. 25 000 total nuclei from this mixture were emulsified in barcoded gel beads using a Chromium Single Cell Controller (10x Genomics). NextGEM Single Cell 3’ Kit V3.1 (10x Genomics) was used for library preparation. A Novaseq 6000 system (Illumina) with a NovaSeq S1 flow-cell was used for paired-end 3’ sequencing of cDNA reads until a total depth of 3.5×10^8^ was achieved.

In parallel, DNA from cells of each genotype was isolated using the GenElute Mammalian Genomic DNA Miniprep Kit Protocol (Merck) and sequenced using the BovineSNP50 v3 DNA Analysis Bead Chip (Illumina) to generate VCF files for each genotype.

#### Data processing and demultiplexing

Demultiplexing based on genotype was used to distinguish nuclei from the different timepoints (0, 24, 48, 72 and 96 h), where each timepoint consisted of two unique genotypes. Demultiplexed FASTQ files were generated from BCL files using the mkfastq function (CellRanger 6.0.1). Using CellRanger’s count function, sequencing reads were mapped to the Bos Taurus genome (build ARS-UCD1.2), annotated (Ensembl release 101) and filtered with default parameters. All reads, including those mapping to introns, were used for subsequent analysis.

The gene expression matrix was obtained by applying default filtering parameters of CellRanger. The previously generated VCF files were used to assign genotype-probabilities to each barcode using demuxlet^66^.

#### General quality control, normalisation, and clustering

In total, we identified 18021 nuclei, of which 4999 were unambiguously categorised as singlets. Low-quality nuclei (beyond three median absolute deviations with respect to expressed genes, total counts or percentage mitochondrial genes) were removed, leaving 4618 nuclei for downstream analysis. These nuclei were normalised using the sctransform function of Seurat with default parameters and clustered by FindNeighbors() and FindClusters() using the first 50 principal components and a resolution of 0.2^67,68^. Of the seven clusters, three were removed due to low quality based on covered SNPs, UMIs per SNP and median read count (clusters 2, 4 and 5) and one because its expression profile indicated contaminating fibro-adipogenic progenitor cells (FAPs; expressing *ITGA5* and lacking *ITGA7*, cluster 6), leaving a total of 3692 nuclei. Due to the lack of spatial information in snRNA-seq data, it was not possible to subdivide the ‘Differentiating’ nuclei into ‘Committed’ and ‘Myotube’ populations in the same fashion as the immunofluorescence data. Cell cycle phases were assigned with the CellCycleScoring() function in Seurat, using the packages’ cc.genes.2019 data set.

#### Dimensionality reduction and differential gene expression analysis

Uniform manifold approximation and projection (UMAP) was used to reduce dimensionality based on the first 50 principal components with 50 neighbouring points of all timepoints collated.

Differentially expressed (DE) genes were computed for the three remaining clusters using FindAllMarkers() with the SCT-normalised assay, a log_2_ fold-change (log-FC) threshold of 0.25 and a false-discovery rate (FDR) of 0.05. DE genes were used for assigning phenotypes to the clusters and for generating GO terms (biological processes, 2023) using enrichR^69^.

### Protein analysis

For 2D samples (Fig. 3d), protein was extracted using RIPA buffer (sc-24948, Santa Cruz Biotechnology). For 3D samples (Figs. 5f, g), protein was extracted using SDS buffer (5% SDS, 50 mM triethylammonium bicarbonate (TEABC; T7408, Sigma)) in combination with mechanical tissue disruption. Total protein was measured using a micro BCA kit (23235, Pierce). Protein samples were boiled in Laemmli buffer (5 min), separated by SDS-PAGE and transferred to PVDF membranes. Equal protein loading was confirmed using Ponceau S Staining (P7170, Sigma). Blots were stained using indicated antibodies (Supplementary Table 2), developed using SuperSignal West Femto Maximum Sensitivity Substrate (Thermo Scientific) and visualised on an Azure 600 chemiluminescence imager (Azure Biosystems). For Fig. 5g, protein samples were extracted from bovine semimembranosus muscle tissue and diluted using SDS sample buffer as indicated.

### Live cell imaging

Live cell imaging (Supplementary Video 1) was carried out using a phase holographic microscope (Phase Focus Ltd) with a 20x objective, 0.9 NA and a lateral resolution of 0.4 µm. Cells were differentiated in 2D culture as described above, and imaged consecutively for 96 h with a frame rate of 3 frames per hour.

### Alginate BAM fabrication

Low molecular weight alginate (L3, Kimica) was purified using a tangential flow filtration system (Sartorius) and subsequently coupled with GGGRGDSP peptide (Biomatik) using DMTMM. This RGD-functionalised alginate was dissolved to 1 wt% and mixed in a 1:1 ratio with SC suspension (5×10^7^ cells ml^-1^) in SFDM v1. Gel mixtures were extruded into 100 mM CaCl_2_ solution for 5 min for crosslinking. Subsequently, BAMs were washed and incubated in SFDM v1. After 48 h, SFDM v2 compounds were added carefully into the medium. After 72 h, medium was exchanged for SFDM v1 for all BAM conditions. BAMs were harvested for protein quantification and confocal microscopy after 168 h. For confocal, samples were fixed, permeabilised and stained as described above for 2D samples, using the indicated antibodies or reagents (Supplementary Table 2). Imaging was performed using a confocal microscope (TCS SP8, Leica Microsystems) with 10x/1.00 objective and 5 μm Z-steps. Presented images are maximum intensity projections.

### Statistical analyses

Statistical significance was assessed using Prism v.9.3.1 (GraphPad). One-way analysis of variance (Figs. 3a, 3c, 4d, 4e and Supplementary Figs. 4a and 6b), two-way analysis of variance (Fig. 4c and Supplementary Figs. 5c, e and f) and t-test (Figs. 5b, 5c, 5d and Supplementary Fig. 5b) were performed to determine statistical significance. Adjusted *p*-values used throughout the figures were: \**p*<0.01, \*\**p*<0.001, \*\*\**p*<0.0001.

### Data availability

snRNA-seq data has been deposited to the GEO (accession number GSE240556). Further data supporting the findings of this study are available from the authors on request.

### Code availability

Code for snRNA-seq analysis and scripts for image analysis are available from the authors on request.

## Acknowledgements

We would like to thank Latifa Karim, Wouter Coppieters, Alice Mayer and Manon Deckers (GIGA institute, ULiège) for support in the acquisition and analysis of snRNA-seq data. We also acknowledge Benjamin Bouchet and Arthur Bister (both Mosa Meat B.V.) for advice on immunofluorescent stainings, André Pötgens (Mosa Meat B.V.) for help with protein measurements and Jessica Rickman (Phase Focus Ltd) for assistance with phase holographic live microscopy. We thank Jessy Chen (Mosa Meat B.V.), Tamar Eigler-Hirsh and Meital Nuriel-Ohayon (both ProFuse Technology) for helpful discussions during this research.

## Author contributions

LM, LS, MF, TM, DR, AK, TvH, SS and JEF performed experiments and analysis. AD, MJP and JEF supervised the study. LM and JEF wrote the manuscript with input from all authors.

