## Supplementary Material for "Optimisation of cell fate determination for cultured muscle differentiation"

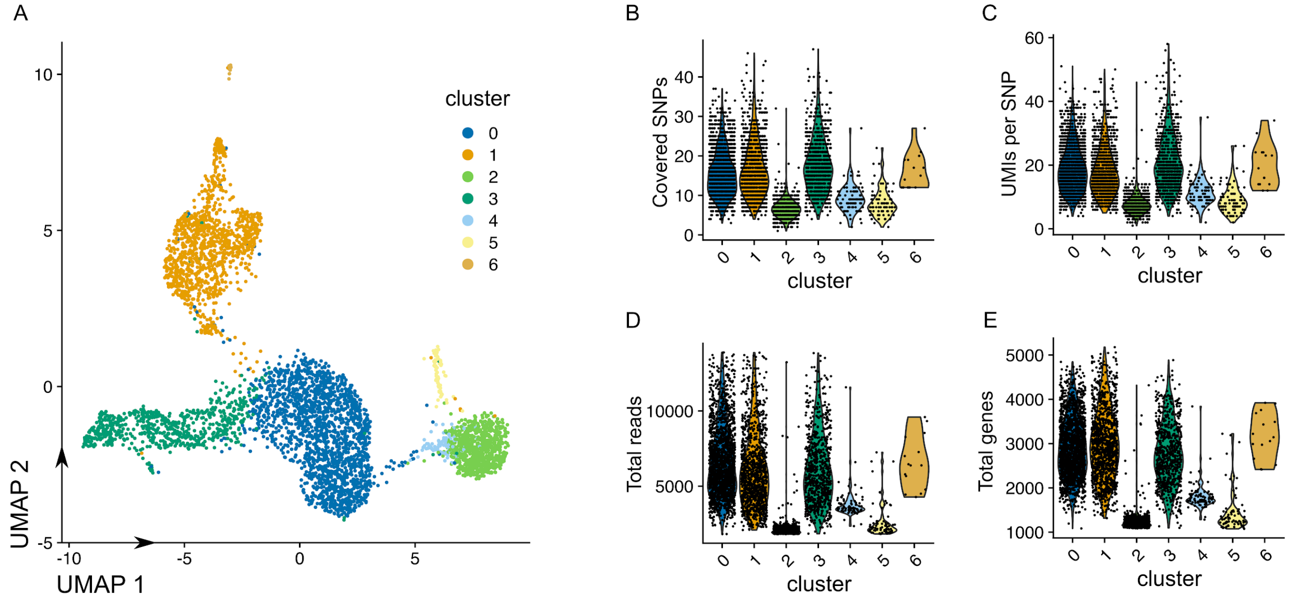


**Supplementary Figure 1: Single-nuclei RNA-sequencing quality control (related to Figure 1)**

A: UMAP of nuclei from all five timepoints prior to quality control, where nuclei are coloured according to clusters identified through shared nearest neighbour (SNN) modularity optimisation.

B: Number of single-nucleotide polymorphisms (SNPs) identified per nucleus, which were used to assign genetic background, where nuclei are grouped and coloured by identified cluster.

C: Average number of unique molecular identifiers (UMIs) overlapping a SNP per nucleus for each cluster.

D: Total number of counts (reads) per nucleus for each cluster.

E: Total number of features (genes) per nucleus for each cluster.

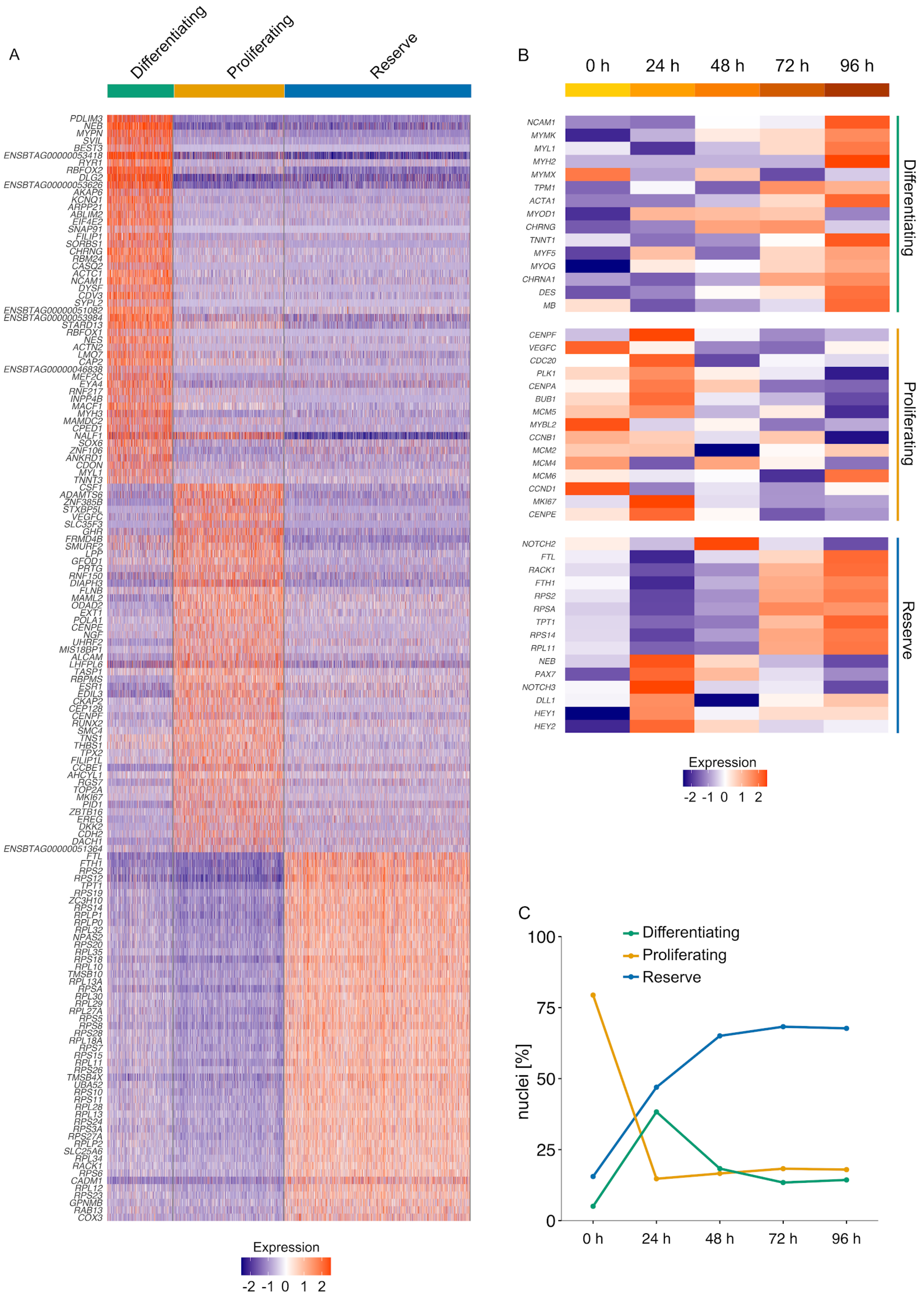
A
Differentiating
Proliferating
Reserve
PDLIM3
NEB
MYPN
SVIL
BEST3
ENSBTAG00000053418
RYR1
RBFOX2
DLG2
ENSBTAG00000053626
AKAP6
KCNQ1
ARPP21
ABLIM2
EIF4E2
SNAP91
FILIP1
SORBS1
CHRNG
RBM24
CASQ2
ACTC1
NCAM1
DYSF
CDV3
SYPL2
ENSBTAG00000051082
ENSBTAG00000053984
STARD13
RBFOX1
NES
ACTN2
LMO7
CAP2
ENSBTAG00000046838
MEF2C
EYA4
RNF217
INPP4B
MACF1
MYH3
MAMDC2
CPED1
NALF1
SOX6
ZNF106
ANKRD1
CDON
MYL1
TNNT3
CSF1
ADAMTS6
ZNF385B
STXBP5L
VEGFC
SLC35F3
GHR
FRMD4B
SMURF2
LPP
GFOD1
PRTG
RNF150
DIAPH3
FLNB
MAML2
ODAD2
EXT1
POLA1
CENPE
NGF
UHRF2
MIS18BP1
ALCAM
LHFPL6
TASP1
RBPMS
ESR1
EDIL3
CKAP2
CEP128
CENPF
RUNX2
SMC4
TNS1
THBS1
TPX2
FILIP1L
CCBE1
AHCYL1
RGS7
TOP2A
MKI67
PID1
ZBTB16
EREG
DKK2
CDH2
DACH1
ENSBTAG00000051364
FTL
FTH1
RPS2
RPS12
TPT1
RPS19
ZC3H10
RPS14
RPLP1
RPLP0
RPL32
NPAS2
RPS20
RPL35
RPS18
RPL10
TMSB10
RPL13A
RPSA
RPL30
RPL29
RPL27A
RPS5
RPS8
RPS28
RPL18A
RPS7
RPS15
RPL11
RPS26
TMSB4X
UBA52
RPS10
RPS11
RPL28
RPL13
RPS24
RPS3A
RPS27A
RPLP2
SLC25A6
RPL34
RACK1
RPS6
CADM1
RPL12
RPS23
GPNMB
RAB13
COX3
Expression
-2 -1 0 1 2
B
0 h
24 h
48 h
72 h
96 h
NCAM1
MYMK
MYL1
MYH2
MYMX
TPM1
ACTA1
MYOD1
CHRNG
TNNT1
MYF5
MYOG
CHRNA1
DES
MB
Differentiating
CENPF
VEGFC
CDC20
PLK1
CENPA
BUB1
MCM5
MYBL2
CCNB1
MCM2
MCM4
MCM6
CCND1
MKI67
CENPE
Proliferating
NOTCH2
FTL
RACK1
FTH1
RPS2
RPSA
TPT1
RPS14
RPL11
NEB
PAX7
NOTCH3
DLL1
HEY1
HEY2
Reserve
Expression
-2 -1 0 1 2
C
Differentiating
Proliferating
Reserve
100
75
50
25
0
nuclei [%]
0 h
24 h
48 h
72 h
96 h

**Supplementary Figure 2: Identification and characterisation of SC subpopulations (related to Figure 2)**

A: Heatmap showing normalised expression of 50 most significantly upregulated genes per SC subpopulation.

B: Heatmap showing average normalised expression of 15 most differentially expressed genes between timepoints within each subpopulation, sorted by UPGMA clustering.

C: Quantification of subpopulation sizes over all five timepoints of the snRNA-seq dataset.

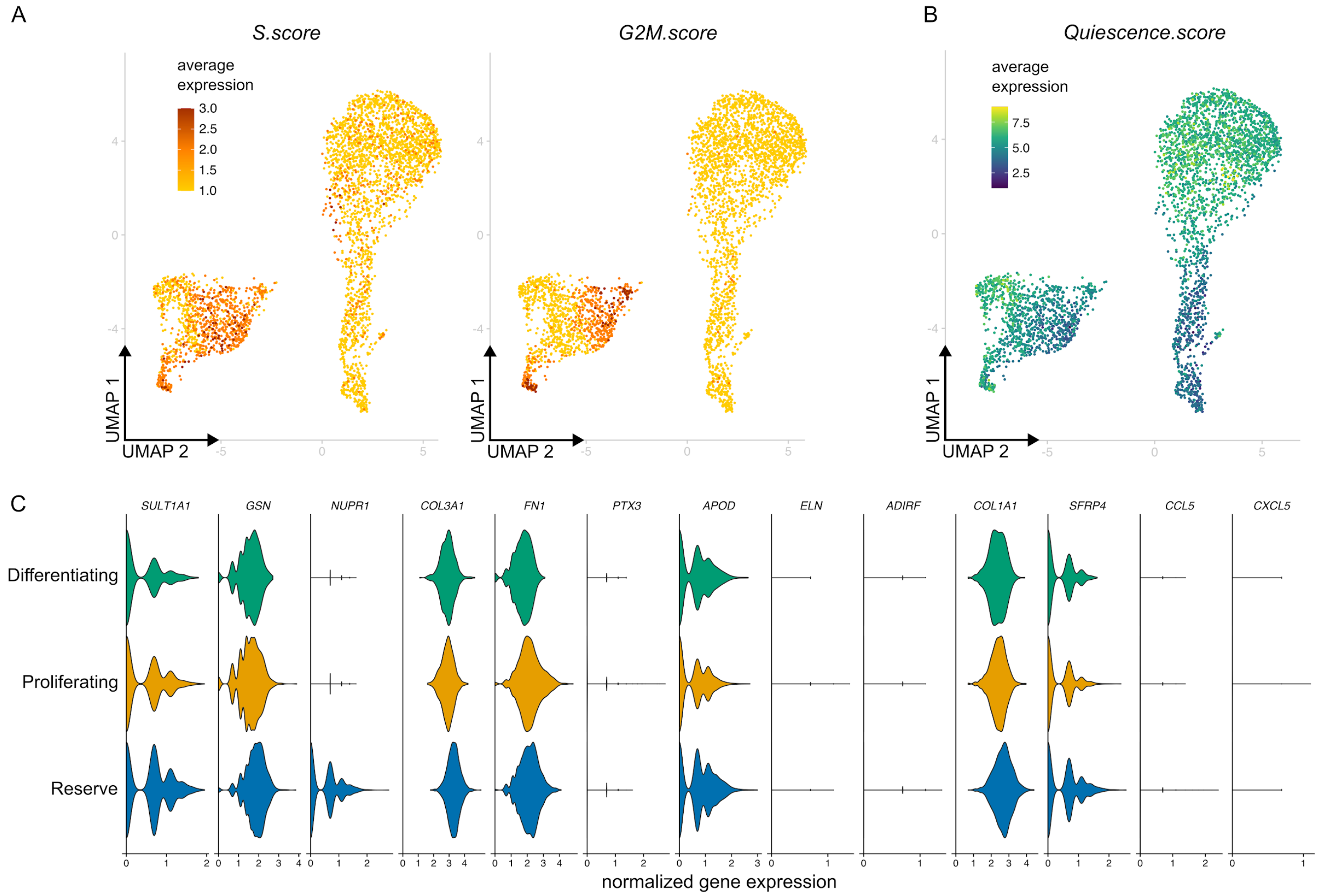


**Supplementary Figure 3: Analysis of cell cycle and quiescence in SC subpopulations (related to Figure 2)**

A: UMAP of nuclei from all five timepoints, coloured by average expression of cell cycle genes related to S (left) or G2M phase (right), as assigned by the CellCycleScoring() function in Seurat.

B: UMAP as in A, coloured by expression of a ‘quiescence signature’ score calculated by averaging over all 597 significantly upregulated genes in a quiescent subpopulation of proliferating SCs (Messmer *et al*., 2023).

C: Violin plots showing normalised expression of 15 most upregulated genes in quiescent SCs (Messmer *et al*., 2023), separated and coloured by subpopulation.

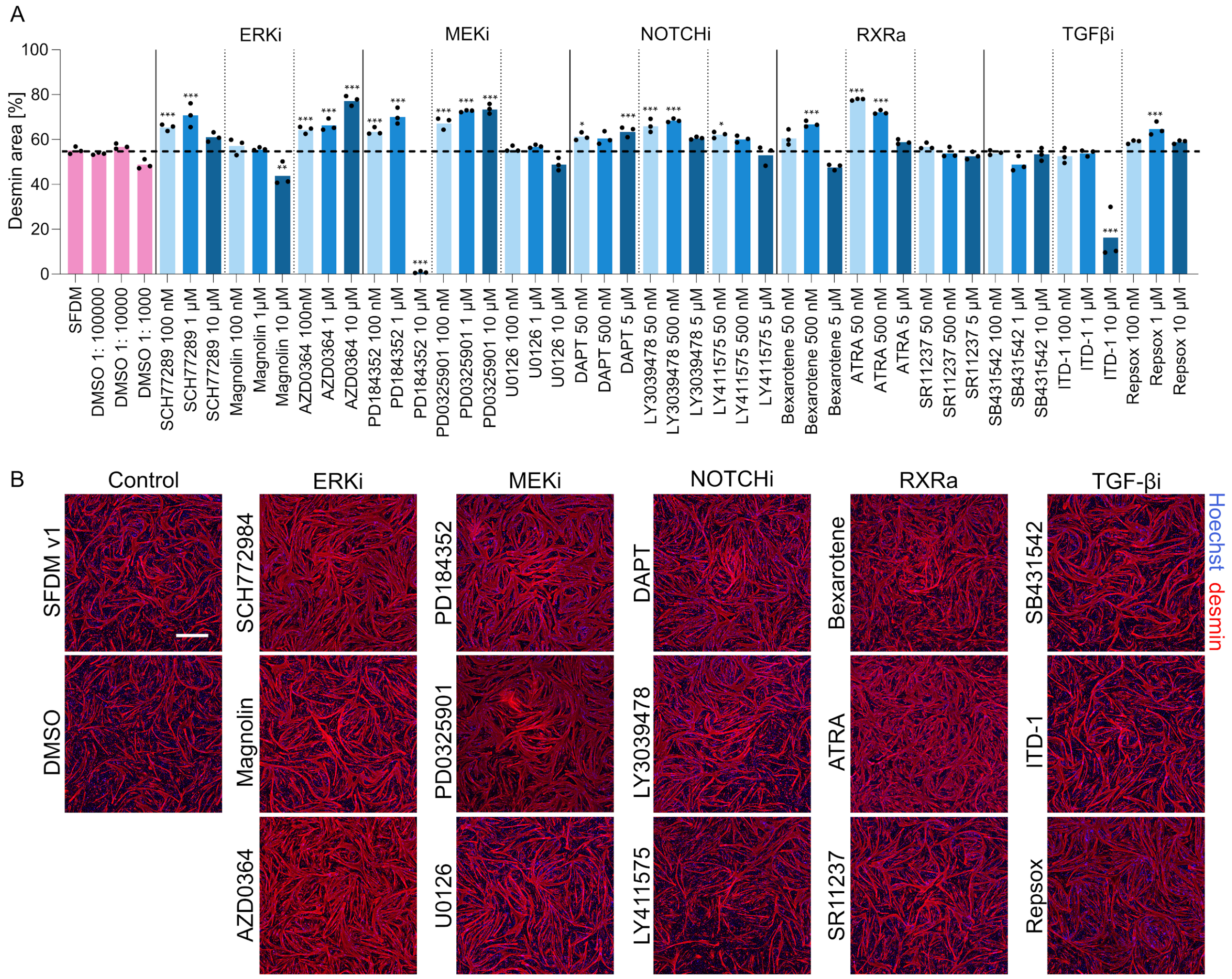


**Supplementary Figure 4: Differentiation compound screening (related to Figure 3)**

A: Mean desmin areas after 72 h differentiation with indicated compounds (targeting MEK/ERK, RXR, NOTCH and TGF-β pathways) tested at the indicated range of concentrations, n = 3.

B: Representative fluorescence images for compound screening shown in Fig. 3a. Red, desmin; blue, Hoechst. Scale bar, 500 μm.

**p*<0.01, ***p*<0.001, ****p*<0.0001.

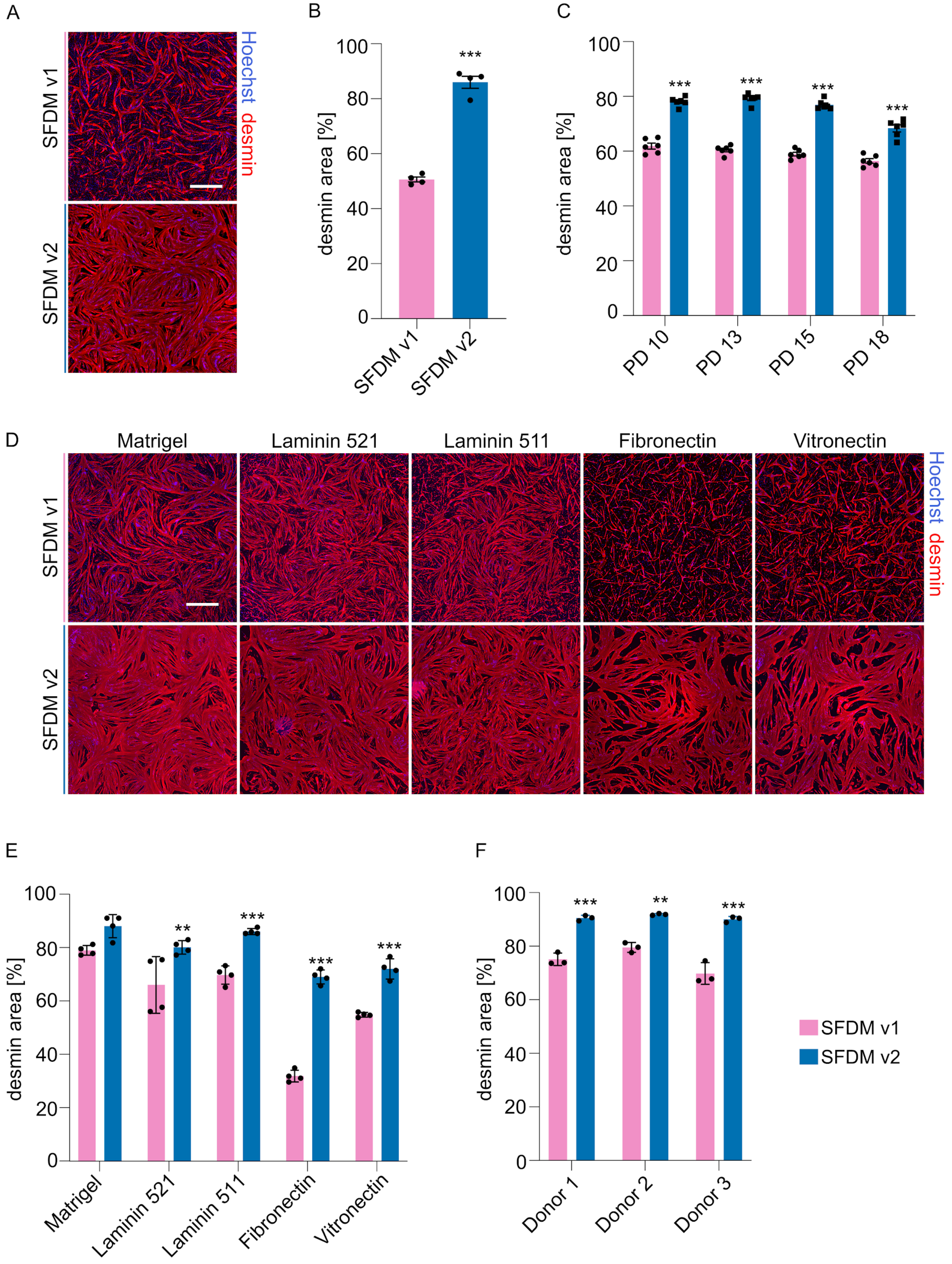
A
SFDM v1
SFDM v2
Hoechst desmin
B
desmin area [%]
100
80
60
40
20
0
***
SFDM v1
SFDM v2
C
desmin area [%]
100
80
60
40
20
0
***
***
***
***
PD 10
PD 13
PD 15
PD 18
D
Matrigel
Laminin 521
Laminin 511
Fibronectin
Vitronectin
SFDM v1
SFDM v2
Hoechst desmin
E
desmin area [%]
100
80
60
40
20
0
**
***
***
***
Matrigel
Laminin 521
Laminin 511
Fibronectin
Vitronectin
F
desmin area [%]
100
80
60
40
20
0
***
**
***
Donor 1
Donor 2
Donor 3
SFDM v1
SFDM v2

**Supplementary Figure 5: Analysis of SFDM v2 for cultured meat bioprocesses (related to Figure 3)**

A: Representative fluorescence images after 72 h myogenic differentiation after proliferation in a 40 L stirred-tank bioreactor. Blue, Hoechst; red, desmin. Scale bar, 500 µm.

B: Mean desmin areas for images in A. Error bars indicate s.d., n = 3.

C: Mean desmin areas after 72 h myogenic differentiation for SCs at different PDs. Error bars show s.d., n = 3.

D: Representative fluorescence images after 72 h of myogenic differentiation on indicated extracellular matrix (ECM) protein coatings. Scale bar, 500 µm.

E: Mean desmin areas for images in D. Error bars show s.d., n = 3.

F: Mean desmin areas after 72 h myogenic differentiation for SCs from three different donor cattle. Error bars show s.d., n = 3.

*$p$<0.01, **$p$<0.001, ***$p$<0.0001.

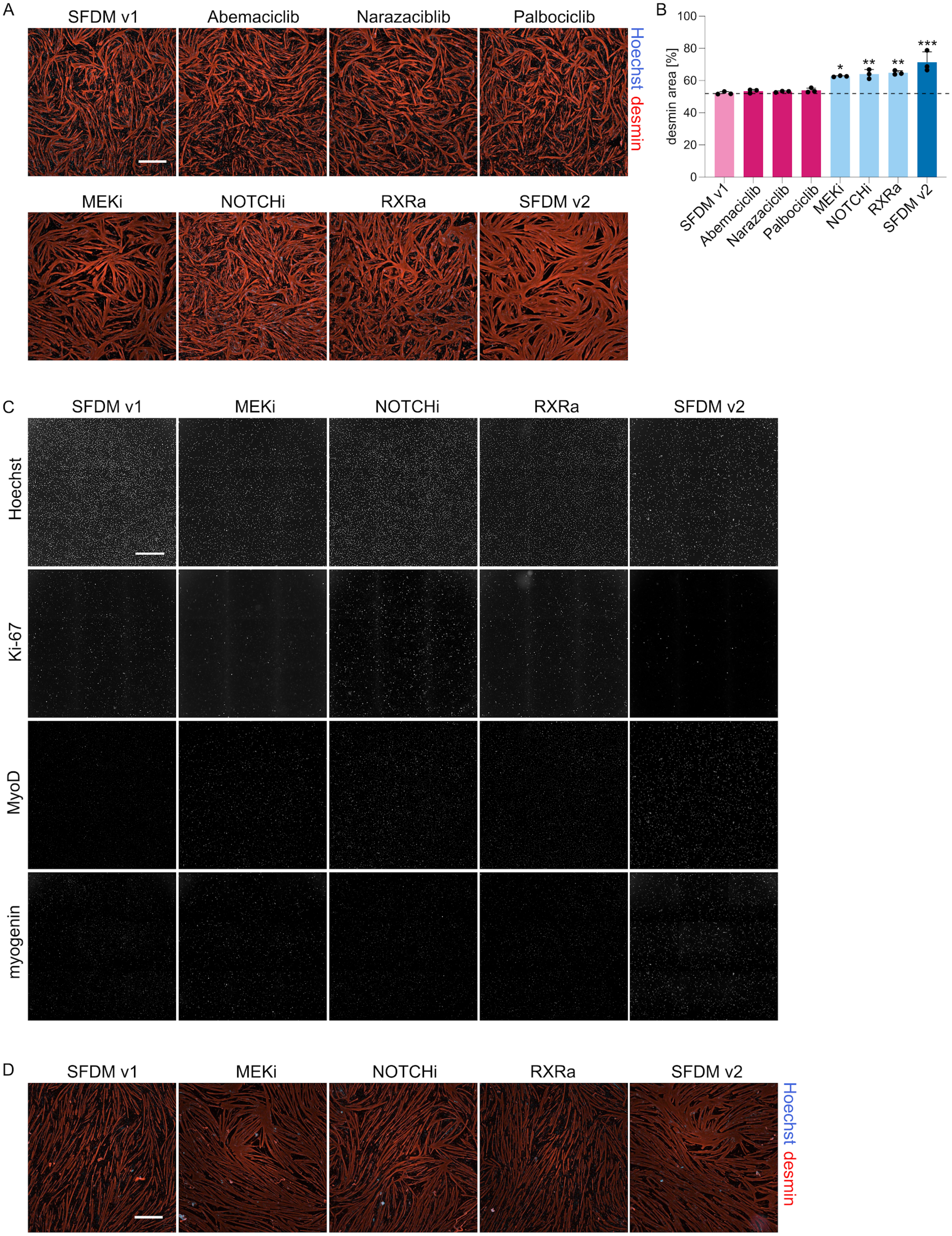
A
SFDM v1
Abemaciclib
Narazaciblib
Palbociclib
MEKi
NOTCHi
RXRa
SFDM v2
Hoechst desmin
B
desmin area [%]
100
80
60
40
20
0
*
**
**
***
SFDM v1
Abemaciclib
Narazaciclib
Palbociclib
MEKi
NOTCHi
RXRa
SFDM v2
C
SFDM v1
MEKi
NOTCHi
RXRa
SFDM v2
Hoechst
Ki-67
MyoD
myogenin
D
SFDM v1
MEKi
NOTCHi
RXRa
SFDM v2
Hoechst desmin

**Supplementary Figure 6: Mechanistic understanding of SFDM v2 components (related to Figure 4)**

A: Representative fluorescence images after 72 h myogenic differentiation with the indicated treatments. Blue, Hoechst; red, desmin. Scale bar, 500 μm.

B: Mean desmin areas for images in A. Error bars indicate s.d., n = 3.

C:  Representative fluorescence images corresponding to Fig. 4d. Target proteins are indicated next to their respective panels. Scale bar, 500 μm.

D: Representative fluorescence images corresponding to Fig. 4e. Blue, Hoechst; red, desmin. Scale bar, 500 μm.

*$p$<0.01, **$p$<0.001, **$p$<0.0001.

**Supplementary Table 1: Medium formulations used in this study**

| Compound | Supplier | Concentration |
|---|---|---|
| **Serum free growth medium (SFGM)** | | |
| DMEM/F-12 | P04-041262B, PAN Biotech | |
| α-linolenic acid | L2376, Sigma Aldrich | 1.0 μg $ml^{-1}$ |
| bFGF-2 | 100-18B, Peprotech | 10 ng $ml^{-1}$ |
| Bovine Serum Albumin (BSA) | A9418, Sigma Aldrich | 5.0 mg $ml^{-1}$ |
| bHGF | 100-39H, Peprotech | 50 ng $ml^{-1}$ |
| Hydrocortisone | H0888, Sigma Aldrich | 0.1 mM |
| Insulin, Transferrin, Selenium, Ethanolamine (ITSE) | 00-101, biogems | 1% |
| GlutaMax | 35050061, ThermoFisher | 2 mM |
| D-glucose | G7021, Sigma Aldrich | 17.7 mM |
| L-ascorbic acid 2-phosphate (Vitamin C) | A8960, Sigma Aldrich | 155 μM |
| Penicillin/Streptomycin/Amphotericin (PSA) | 17-745E, Lonza | 1% |
| T3 | T6397, Sigma Aldrich | 20 ng $ml^{-1}$ |
| **Serum free differentiation medium version 1 (SFDM v1)** | | |
| DMEM | A14430-01, Gibco | |
| EGF-1 | AF-100-15, Peprotech | 10 ng $ml^{-1}$ |
| D-glucose | G7021, Sigma | 5.5 mM |
| GlutaMax | 35050061, ThermoFisher | 2 mM |
| Human Serum Albumin | Rc HA NW20, Richcore Lifesciences | 0.5 mg $ml^{-1}$ |
| ITSE | 00-101, biogems | 2% |
| L-ascorbic acid 2-phosphate (Vitamin C) | A8960, Sigma Aldrich | 40 μM |
| Lysophosphatidic acid (LPA) | L7260, Sigma Aldrich | 1 μM |
| MEM Amino Acids Solution | 11130-051, ThermoFisher | 0.50% |
| $NaHCO_3$ | P2256, Sigma Aldrich | 6.5 mM |
| PSA | 17-745E, Lonza | 1% |
| Soy hydrolysates | 58903C, Merck | 1% |
| Sodium l-lactate | 71718, Sigma | 10 mM |
| Sodium pyruvate | P2256, Sigma Aldrich | 0.5 mM |
| **Serum free differentiation medium version 2 (SFDM v2)** | | |
| SFDM v1 + | | |
| ATRA (trans-retinoic acid) | 554720, Millipore | 500 nM |
| DAPT, gamma-Secretase inhibitor | ab120622, Abcam | 5 μM |
| PD0325901 | S1036, Selleckchem | 1 μM |

### Supplementary Table 2: Antibodies used in this study

| Target | Colour | Supplier | Reference | Dilution | Application |
|---|---|---|---|---|---|
| $\alpha$-actin-1 | - | Abcam | ab184705 | 1:5,000 | Western Blot |
| actinin | - | Sigma-Aldrich | A7811 | 1:1,600 (IF); 1:2,500 (WB) | IF Western Blot |
| f-actin | Atto550 | Sigma-Aldrich | 19083 | 1:300 | IF |
| desmin (rabbit) | - | Abcam | ab227651 | 1:400 | IF |
| desmin (goat) | - | Abcam | ab80503 | 1:100 | IF |
| Ki-67 | - | Miltenyi Biotech | 130-108-060 | 1:100 | IF |
| MyoD | - | LifeSpan BioSciences | LS-C88010-100 | 1:100 | IF |
| myogenin | - | Abcam | ab219998 | 1:100 | IF |
| myoglobin | - | Abcam | ab231725 | 1:2,500 | Western Blot |
| myosin | - | Abcam | ab51263 | 1:2,000 | IF |
| myosin | - | Abcam | ab11083 | 1:5,000 | Western Blot |
| Pax7 | - | Developmental Studies Hybridoma Bank | Cat# PAX7 | 1:100 | IF |
| tubulin | - | Abcam | ab4074 | 1:10,000 | Western Blot |
| goat | AF546 | Invitrogen | A-11056 | 1:300 | IF |
| human | DyLight488 | Invitrogen | SA5-10126 | 1:300 | IF |
| mouse | AF647 | Invitrogen | A32787 | 1:300 | IF |
| mouse | AF488 | Invitrogen | A-11001 | 1:1,000 | IF |
| mouse | AF633 | Thermo Fisher Scientific | A21050 | 1:500 | IF |
| mouse | DyLight650 | biotechne | NBP1-75147C | 1:300 | IF |
| mouse | HRP | DAKO | P0447 | 1:2,000 | Western Blot |
| rabbit | AF488 | Thermo Fisher Scientific | A21206 | 1:300 | IF |
| rabbit | DyLight550 | Thermo Fisher Scientific | SA5-10033 | 1:250 | IF |
| rabbit | AF594 | Thermo Fisher Scientific | R37119 | 1:300 | IF |
| rabbit | HRP | Abcam | ab6721 | 1:10,000 | Western Blot |

**Supplementary Table 3: Small molecule compound screening**

| Compound | Supplier | Concentration | Pathway |
|---|---|---|---|
| Abemaciclib | MedChemExpress | 100 nM | CDKi |
| Narazaciclib | MedChemExpress | 100 nM | CDKi |
| Palbociclib | MedChemExpress | 100 nM | CDKi |
| SCH772984 | Selleckchem | 1 μM | ERKi |
| Magnolin | Selleckchem | 1 μM | ERKi |
| AZD0364 | Selleckchem | 1 μM | ERKi |
| PD184352 | Selleckchem | 1 μM | MEKi |
| PD0325901 | Selleckchem | 1 μM | MEKi |
| U0126-EtOH | Selleckchem | 1 μM | MEKi |
| DAPT | Abcam | 5 μM | NOTCHi |
| LY3039478 | Selleckchem | 5 μM | NOTCHi |
| LY411575 | Miltenyi Biotec | 5 μM | NOTCHi |
| Bexarotene | Selleckchem | 500 nM | RXRa |
| All-trans-retinoic acid (ATRA) | Millipore | 500 nM | RXRa |
| SR11237 | Sigma Aldrich | 500 nM | RXRa |
| SB431542 | Selleckchem | 1 μM | TGF-βi |
| ITD-1 | MedChemExpress | 1 μM | TGF-βi |
| Repsox | Selleckchem | 1 μM | TGF-βi |

**Supplementary Video 1: Live imaging of differentiating SCs (related to Figure 3)**

Representative phase holographic microscopy videos of 2D myogenic differentiation over 96 h with indicated treatments. Scale bar, 500 μm.
